# Characterization of Intellectual disability and Autism comorbidity through gene panel sequencing

**DOI:** 10.1101/545772

**Authors:** Maria Cristina Aspromonte, Mariagrazia Bellini, Alessandra Gasparini, Marco Carraro, Elisa Bettella, Roberta Polli, Federica Cesca, Stefania Bigoni, Stefania Boni, Ombretta Carlet, Susanna Negrin, Isabella Mammi, Donatella Milani, Angela Peron, Stefano Sartori, Irene Toldo, Fiorenza Soli, Licia Turolla, Franco Stanzial, Francesco Benedicenti, Cristina Marino-Buslje, Silvio C.E. Tosatto, Alessandra Murgia, Emanuela Leonardi

## Abstract

Intellectual disability (ID) and autism spectrum disorder (ASD) are clinically and genetically heterogeneous diseases. Recent whole exome sequencing studies indicated that genes associated with different neurological diseases are shared across disorders and converge on common functional pathways. Using the Ion Torrent platform, we developed a low-cost next generation sequencing (NGS) gene panel that has been transferred into clinical practice, replacing single disease gene analyses for the early diagnosis of individuals with ID/ASD. The gene panel was designed using an innovative *in silico* approach based on disease networks and mining data from public resources to score disease-gene associations. We analyzed 150 unrelated individuals with ID and/or ASD and a confident diagnosis has been reached in 26 cases (17%). Likely pathogenic mutations have been identified in another 15 patients, reaching a total diagnostic yield of 27%. Our data also support the pathogenic role of genes recently proposed to be involved in ASD. Although many of the identified variants need further investigation to be considered disease-causing, our results indicate the efficiency of the targeted gene panel on the identification of novel and rare variants in patients with ID and ASD.

## INTRODUCTION

Neuro-developmental disorders (NDDs) are common conditions including clinically diverse and genetically heterogeneous diseases. Intellectual disability (ID) is the most common NDD disorder, with a prevalence varying between 0.5 to 3% in general population, depending on patient and parent age, or the measure of intellectual quotient used (Leonard et al., 2011). ID is characterized by deficits in both intellectual and adaptive functioning that first manifest during early childhood. Children with intellectual disability (ID) exhibit increased risk to present potential co-occurring developmental conditions, such as autism spectrum disorders (ASDs) (28%), epilepsy (22.2%), stereotypic movement disorders (25%), and motor disorders, which substantially affect daily living and well-being (Almuhtaseb et al., 2014; Jensen and Girirajan, 2017; Kazeminasab et al., 2018). ASD in particular is characterized by deficits in social communication and interactions, as well as by repetitive behaviors and restrictive interests, is associated with poorer psychosocial and family related outcomes than ID alone (Totsika et al., 2011). ASD as well as epilepsy commonly coexist in specific neurodevelopmental disorders with ID, such as Fragile-X and Rett syndromes, or in phenotypes associated with specific copy number variations (CNVs) and single gene mutations. These make the differential diagnosis among these disorders extremely difficult based only on clinical features. Furthermore, it seems that patients affected by one of these disorders have high risk to develop other comorbid NDDs. Exome sequencing studies of family trios with ID, ASD, and epilepsy have revealed a significant excess of *de novo* mutations in probands, when compared to the normal population, and yielded a rich source of candidate genes contributing to these neurodevelopmental defects (Neale et al., 2012; Epi4K Consortium et al., 2013; Fromer et al., 2014). It has been estimated that mutations in more than 1,000 different genes might cause ID (Chiurazzi and Pirozzi, 2016). Both common and rare genetic variants in up to 1,000 genes have been linked to increased ASD risk (SFARI database, https://gene.sfari.org/). However, significant numbers of genes harboring *de novo* mutations are shared across different neurodevelopmental or neuropsychiatric disorders (Vissers et al., 2010; Cukier et al., 2014). Many genes have already been shown to cause both ID and/or ASD, including *PTCHD1*, *SHANK3, NLGN4, NRXN1, CNTNAP2, UBE3A, FMR1, MECP2*, and others (Harripaul et al., 2017). Despite the apparent distinct pathogenesis for these disorders, analysis of network connectivity of the candidate genes revealed that many rare genetic alterations converge on a few key biological pathways (Vissers et al., 2010; Krumm et al., 2014). In the case of ASD, diverse integrative systems biology approaches highlighted how disease genes cluster together in networks enriched in synaptic function, neuronal signaling, channel activity, and chromatin remodeling (Gilman et al., 2012; O’Roak et al., 2012; Pinto et al., 2014). Accordingly, many ASD genes are synaptic proteins, chromatin remodelers, or FMRP targets, i.e. genes encoding transcripts that bind to FMRP (Iossifov et al., 2012). Alterations of the same classes of protein functions and biological processes involved in neuronal development, such as the mammalian target of rapamycin (mTOR) pathways, GABA receptor function or glutamate NMDA receptor function, have been also found implicated in intellectual disability, epilepsy, and schizophrenia (Endele et al., 2010; Gilman et al., 2011; Paoletti et al., 2013; Cristino et al., 2014; Krumm et al., 2014; Reijnders et al., 2017). The multiple genes and molecular pathways shared by ID, ASD and other developmental or psychiatric disorders indicate a common origin that explains the co-occurrence of these conditions (Barabási et al., 2011; Cukier et al., 2014). Based on the hypothesis that common functional pathways explain comorbidity between diverse NDDs disorders, we developed an efficient and cost-effective amplicon-based multigene panel to assess the pathogenic role of genes involved in ID and ASDs comorbidity. The 74-gene panel was designed using an innovative *in silico* approach based on disease networks and mining data from public resources to score disease-gene association. Here, we present the genetic findings after applying this panel to 150 individuals from our cohort of individuals with ID and/or ASD, most of them were negative for array-CGH (aCGH), Fragile-X test and other specific genetic analyses (*MECP2*, *CDKL5*, *UBE3A*, chr15q methylation test, etc.). We adopted a manual prioritization procedure based on expert knowledge related to the disease phenotype and gene functions, which allowed detecting a causative or likely pathogenic variant in 27% of these patients. We describe diagnosed cases that highlight the critical steps of variant interpretation, in the clinical diagnostic context of neurodevelopmental conditions such as ID and ASDs. For each tested individual, we report a clinical description and genetic data from the 74 genes providing a set of genotype – phenotype associations, which can be used to train or test computational methods for prioritization of potential disease-causing variants.

## MATERIALS AND METHODS

### Patient selection

Patients were referred from clinical geneticists of 17 Italian public hospitals with a diagnosis of non-specific neurodevelopmental disorder. Clinical data were collected with a standardized clinical record describing clinical and family history, clinical phenotype (auxological parameters, neurological development, physical features, and behavioral profile), and presence of associated disorders. Data from neurophysiological profiles, electroencephalograms (EEG) and brain magnetic resonance imaging (MRI) were also collected. Table 1 summarizes the clinical data of the patients. Written informed consent was obtained from the patient’s parents or legal representative. This study was approved by the Local Ethics Committee, University-Hospital of Padova, Italy.

**Table 1:** Description of the cohort of 150 individuals enrolled for the study of ID/ASD comorbidity.

| Features | Patients (n=150) | Yield with causative mutations | Yield with causative or putative mutations |
| --- | --- | --- | --- |
| **Gender** |  |  |  |
| Female | 58 (39%) | 10 (17.2%) | 17 (29.3%) |
| Male | 92 (61%) | 16 (17.4%) | 24 (26%) |
| Age (at diagnosis) | 2- 42 y.o | 4-27 y.o. | 3-42 y.o. |
| **Familial History** |  |  |  |
| Sporadic | 121 (81%) | 23 (19%) | 31(25.6%) |
| Familial | 29 (19%) | 3 (10.3%) | 10 (34.5%) |
| Sib pair | 5 (3.3%) | 0 | 0 |
| X-linked | 3 (2%) | 1 | 0 |
| **Intellectual Disability** | **146 (97.3%)** | **26 (17.8%)** | **41 (28%)** |
| Mild | 37 (24.6%) | 5 (13.5%) | 8 (21.6%) |
| Moderate | 38 (25.3%) | 8 (21%) | 10 (26.3%) |
| Severe | 33 (22%) | 9 (27.3%) | 12 (36.3%) |
| Not evaluated | 38 (25.3%) | 4 (10.5%) | 11 (28.9%) |
| **Comorbidity** |  |  |  |
| ASD (autistic features) | 93 (62%) | 15 (16.1%) | 25 (26.8%) |
| ASD not reported | 16 (10.6%) | 3 (18.7%) | 5 (31.2%) |
| Epilepsy | 55 (36.6%) | 11 (20%) | 17 (30.9%) |
| Hypotonia | 28 (18.6%) | 6 (21.4%) | 6 (21.4%) |
| Ataxia | 11 (7.3%) | 2 (18%) | 3 (27.2%) |
| Microcephaly | 19 (12%) | 6 (31.5%) | 8 (42.1%) |
| Macrocephaly | 11 (7.3%) | 4 (36.4%) | 4 (36.4%) |
| **Previous investigation** |  |  |  |
| aCGH | 125 (83.3%) | 22 (17.6%) | 35 (28%) |
| Fragile-X | 86 (57.3%) | 10 (11.6%) | 18 (20.9%) |
| EEG anomaly | 56 (37.3%) | 9 (16%) | 17 (30.4%) |
| MRI anomaly | 37 (24.6%) | 9 (24.3%) | 13 (35.1%) |
| Other tests | 57 (38%) | 13 (22.8%) | 18 (31.6%) |

### Gene panel selection

For the construction of an efficient and low-cost gene panel, we selected the most promising ID and/or ASD genes gathering data from public databases (AutismKB http://autismkb.cbi.pku.edu.cn, and SFARI https://sfari.org/resources/sfari-gene, Abrahams et al. 2013), OMIM, and PubMed. Candidate genes were extracted in particular from recent exome sequencing and meta-analysis studies (Supplementary Table S1). We collected a list of 972 genes scored according to recurrence in different sources, annotated for clinical phenotype, gene function, subcellular localization, and interaction with other known causative genes. Separated lists were generated considering ASD or ID association. The extracted information was stored in a dedicated SQL database used in conjunction with the disease network construction. Using data from STRING 9.0 (Franceschini et al., 2013), a disease protein-protein interaction (PPI) network was built starting from 66 high confidence genes (intersection list), shared by both ASD and ID gene lists. Emerging features of the network were assessed by enrichment analysis with Enrich web-server (Kuleshov et al., 2016). The same list was used as training set for Endeavour gene prioritization (https://endeavour.esat.kuleuven.be/) (Tranchevent et al., 2016). Hub direct interactors (i.e. genes with STRING degree score above 0.45) belonging to the top ranking prioritized list, but not included in the intersection, were also included in the most promising candidate gene list. To this list, we also added the top ranked genes associated to ID or ASD only (i.e. genes with at least 5 evidences for ID or ASD) (Supplementary Figure S1). The final panel set resulted in a manually curated list of 74 genes, comprising selected known causative genes, top-ranked genes by gene prioritization, and genes meeting PPI network parameters (Supplementary Table S2).

### Gene Panel Sequencing

Nucleic acids were extracted from blood samples using the Wizard genomic DNA Promega Kit (Promega Corporation). Multiplex, PCR-based primer panels were designed with Ion AmpliSeq™ Designer (Thermo Fisher Scientific) to amplify all exons and flanking regions (10 bp) of the 74 selected genes (Thermo Fisher Scientific). Template preparation and enrichment were performed with the Ion One Touch 2 and Ion One Touch ES System, respectively (Thermo Fisher Scientific). Read alignment to the human genome reference (hg19/GRCh37) and variant calling were performed with the Ion Torrent Suite Software v5.02 (Thermo Fisher Scientific).

### Variant filtering

An in house pipeline was built to create a database of genetic variants identified in our cohort and to annotate them with features provided by ANNOVAR, i.e. allelic frequency (AF) in control cohorts, variant interpretation from InterVar automated, ClinVar report, pathogenicity predictions and conservation scores. Detected variants were ranked for their frequency in the gnomAD (Lek et al., 2016), 1000G (1000 Genomes Project Consortium et al., 2015), and ExAC (Kobayashi et al., 2017) databases, as well as in our in house database of 150 patients. We excluded SNVs found more than twice in our cohort or reported with an AF higher than expected for the disorder, which has been calculated to be < 0.002% and <0.45% for autosomal dominant and recessive genes, respectively (Piton et al., 2013). Variants reported as risk factors for autism in the literature were nevertheless considered for further segregation analysis even if their frequency in the control populations exceeded the ID incidence. Rare variants were ranked for their pathogenicity prediction considering the consensus among twelve computational methods. ANNOVAR provides predictions from SIFT (Sim et al., 2012), the Polyphen-2 (Adzhubei et al., 2013) HDIV and HVAR versions, LRT (Chun and Fay, 2009), Mutation Taster (Schwarz et al., 2014), MutationAssessor (Reva et al., 2011), FATHMM (Shihab et al., 2014); PROVEAN (Choi and Chan, 2015), MetaSVM (Dong et al., 2015), MetaLR (Dong et al., 2015), M-CAP (Jagadeesh et al., 2016), fathmmMKL (Shihab et al., 2013, 2014) and CADD (Kircher et al., 2014). Conservation was evaluated with the scoring schemes GERP++ (Davydov et al., 2010), PhyloP (Pollard et al., 2010) and SiPhy (Garber et al., 2009). Intronic or synonymous variants near the exon-intron junction were also evaluated *in silico* for their impact on splicing using Human Splicing Finder (Desmet et al., 2009). The Integrated Genome Viewer platform (Robinson et al., 2011) has been used to exclude sequencing or alignment errors around selected SNVs.

### Variant validation and functional assays

Selected variants were validated by Sanger sequencing. Segregation analysis was performed in the patient relatives when DNA samples were available. For apparent *de novo* variants, paternity and maternity were confirmed by the inheritance of rare detected variants in parental samples. In other cases pedigree concordance was checked using polymorphic microsatellite markers of chr15q described in (Giardina et al., 2008). For maternally inherited X-linked variants, the X-inactivation pattern of the mother was evaluated on the highly polymorphic androgen receptor (ARA locus) at Xq11-q12, as described in (Bettella et al., 2013). The X‐inactivation was classified as random (ratio < 40:60) or significantly skewed (ratio ≥ 80:20).

Analysis of the transcripts was performed to confirm putative splicing variants. RNA was extracted from patient peripheral blood leukocytes and real-time PCR performed using random primers. cDNA was used as template in nested PCR reactions with specific primers in order to amplify the regions containing the mutation. PCR products were tested on 1.5% agarose gel and sequenced.

### Variant classification

A clinical interpretation of selected variants was first evaluated using the InterVar (Li and Wang, 2017), adjusting the automated classification with our own findings. The American College of Medical Genetics and Genomics (ACMG) criteria (Richards et al., 2015) used to classify the variants are reported for both causative and likely pathogenic variants (Tables 2 and 3). All the identified variants have been submitted to LOVD database.

**Table 2:**
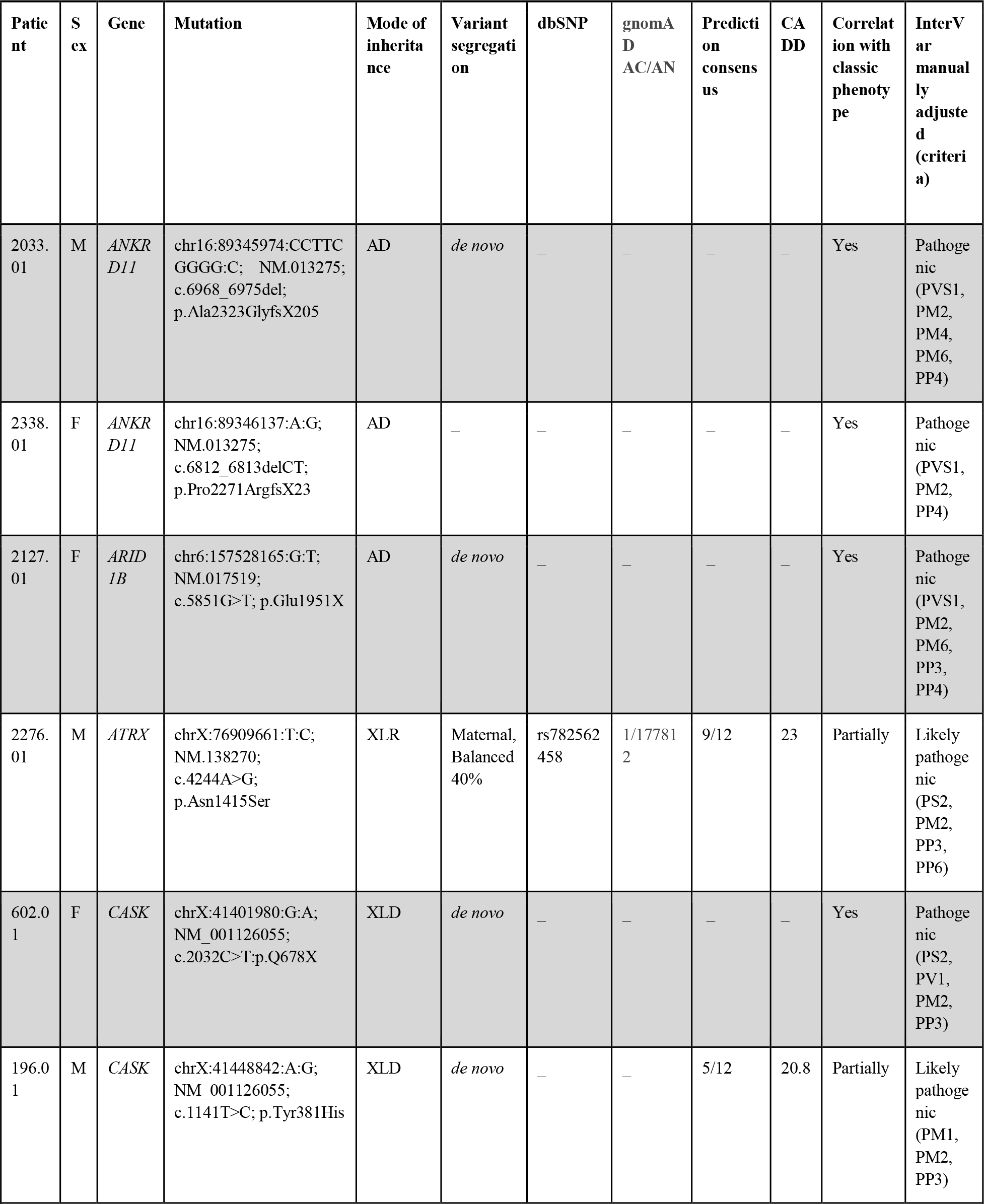
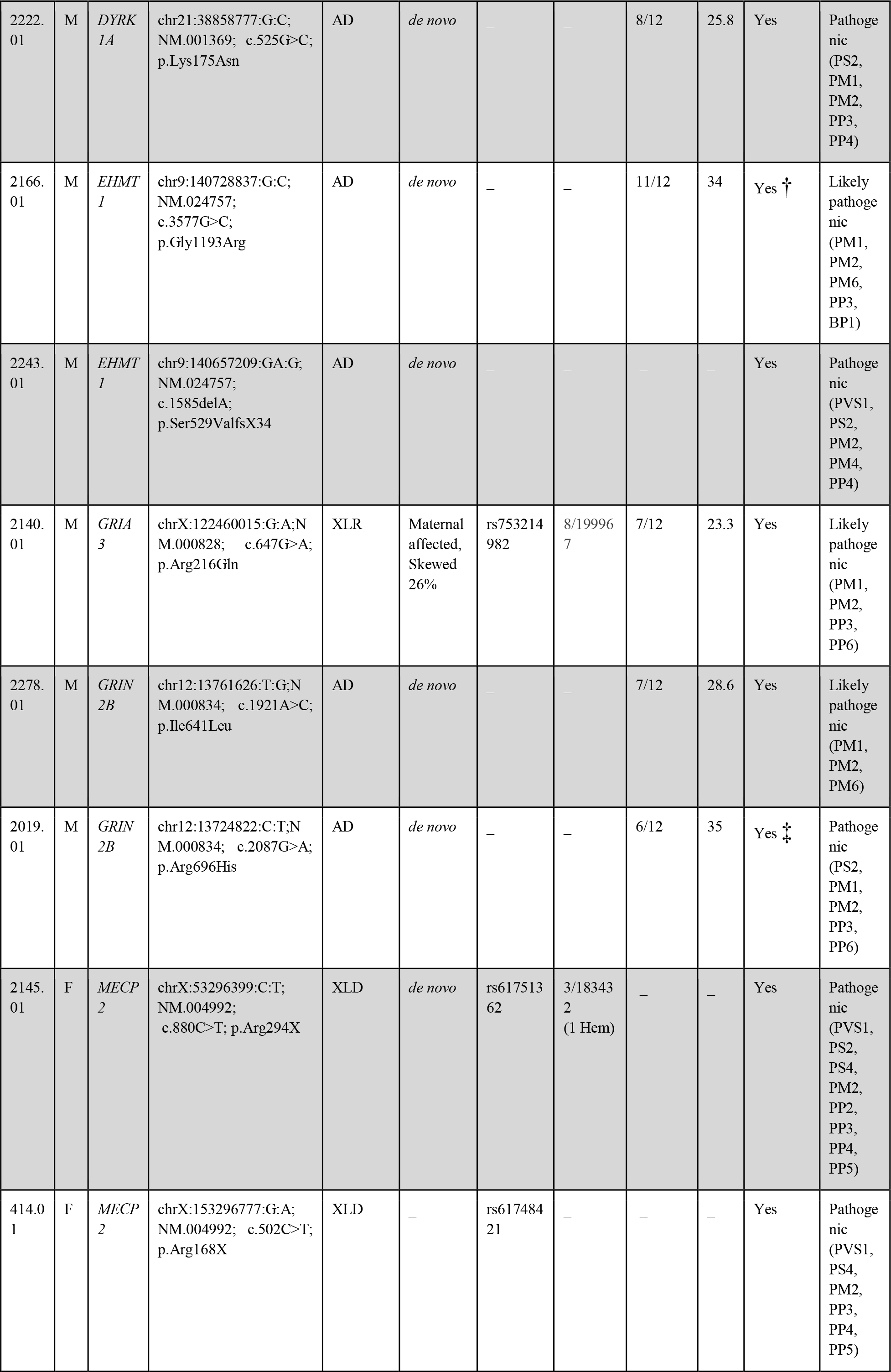
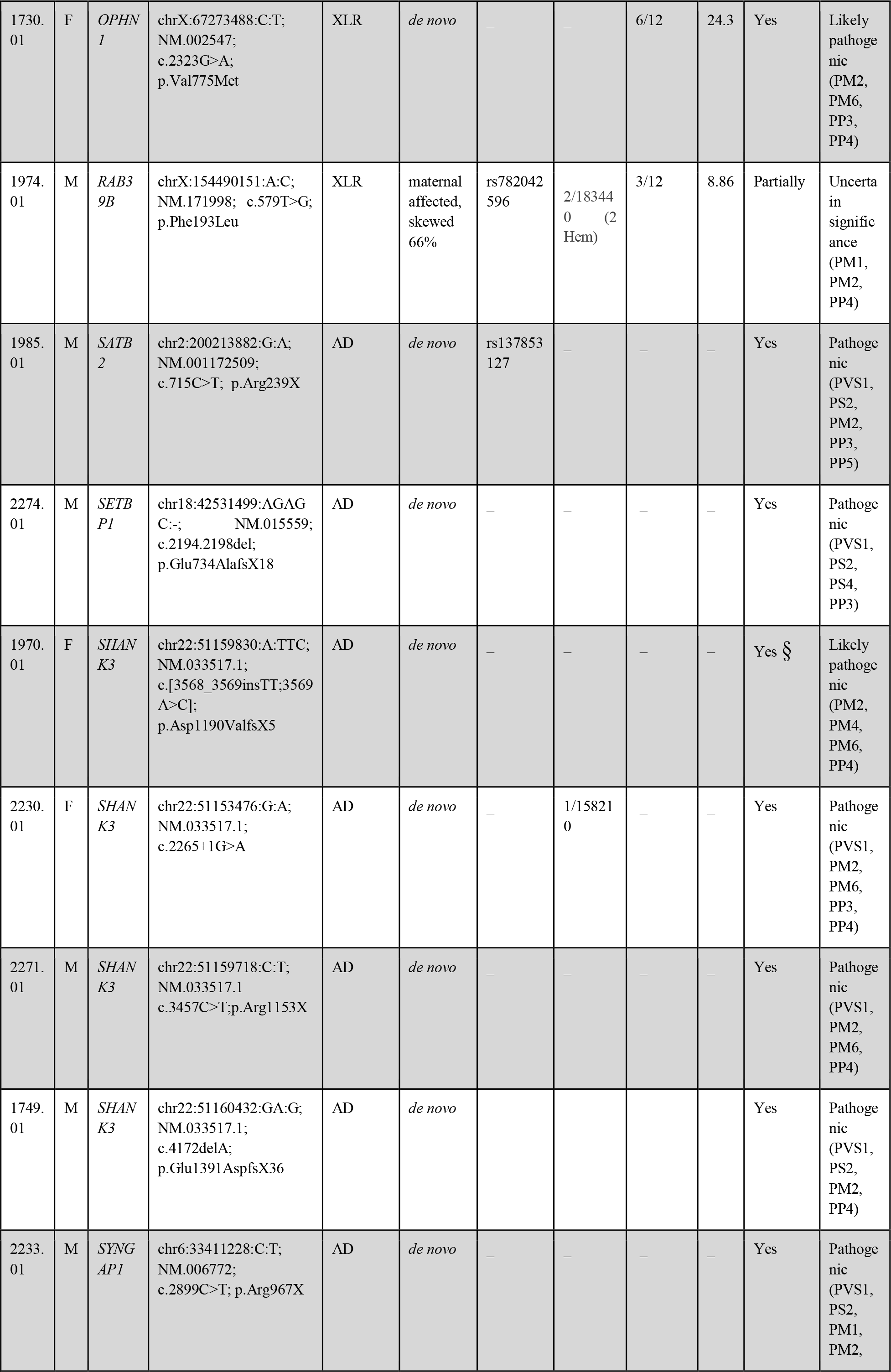
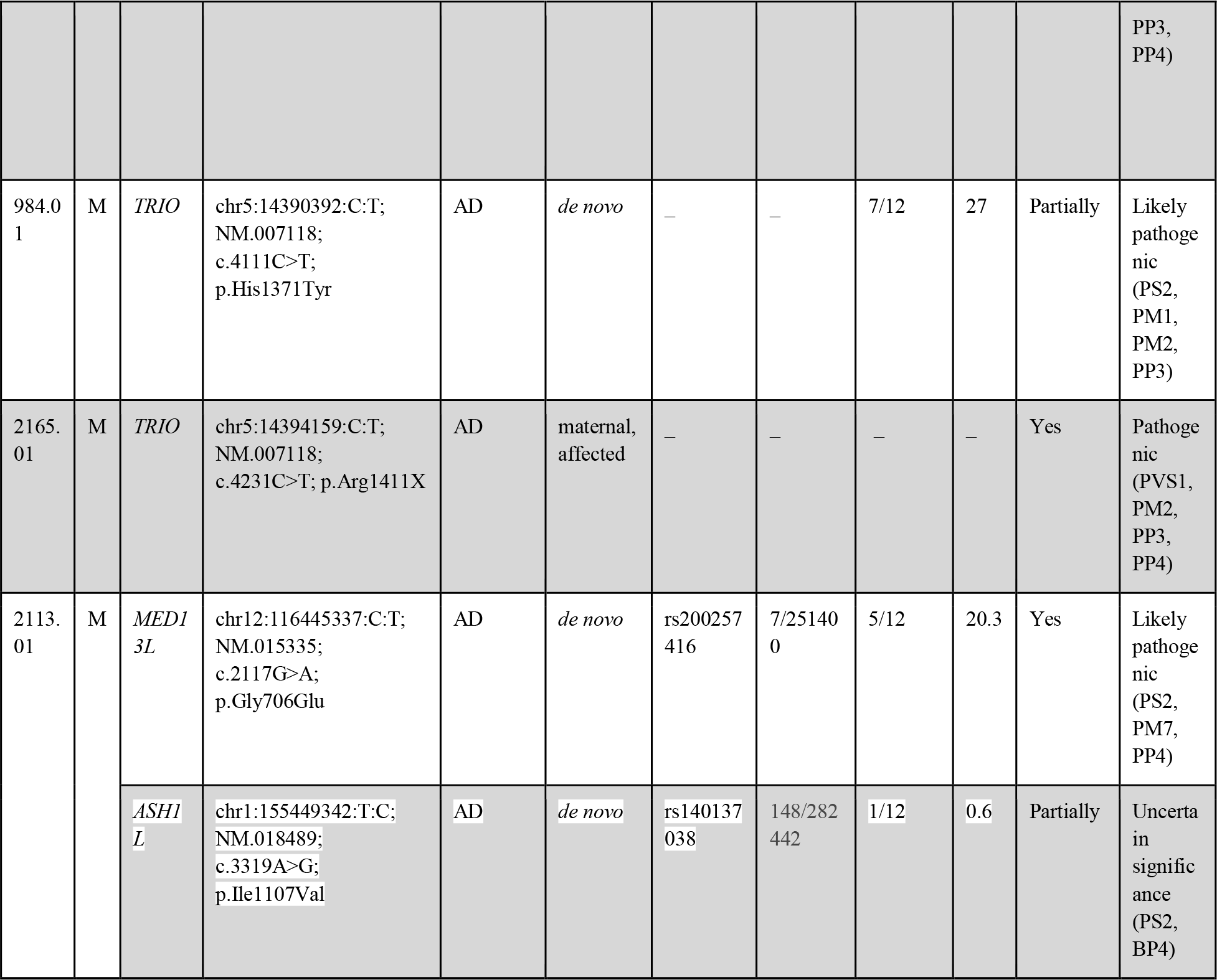
Causative variants found in the 150 patient cohort. For maternally inherited variants, based on X-inactivation analysis, the quote of expressed mutated allele in the mother was reported. Prediction consensus is calculated among the twelve methods provided by ANNOVAR (see Methods section). Variant interpretation based on InterVar: criteria manually adjusted based on our findings (Richards et al., 2015). M, male; F, female; Genomic position on human GRCh37; AD: autosomal dominant; AR: autosomal recessive; XLD: X-linked dominant; XLR: X-linked recessive; † according to the *EHMT1* phenotypic spectrum described in (Blackburn et al., 2017); ‡ This *GRIN2B* missense mutation has been previously found in a female with a phenotype similar to our case (Swanger et al., 2016). § This *SHANK3* splicing mutation has been previously reported by (Li et al., 2018).

**Table 3:**
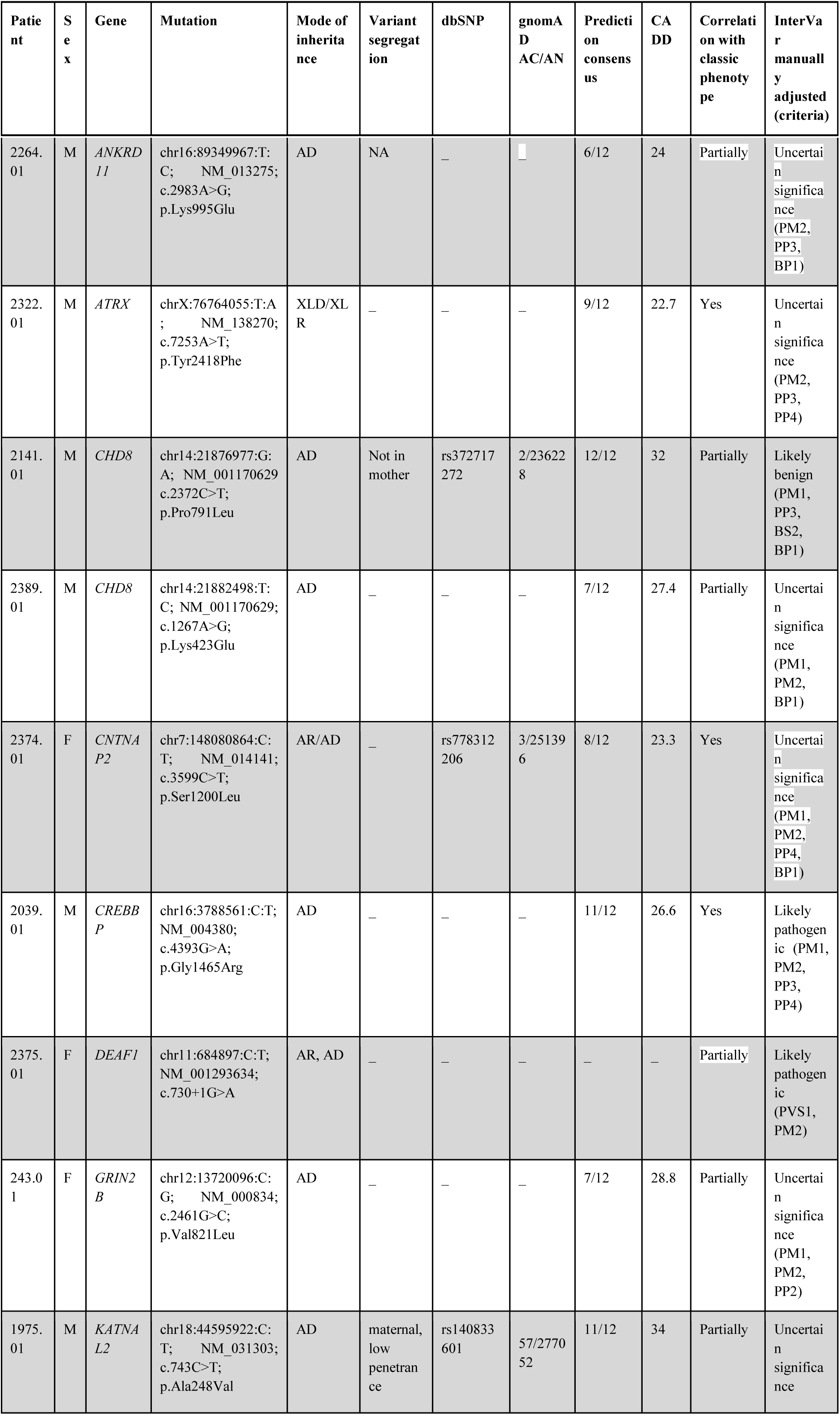
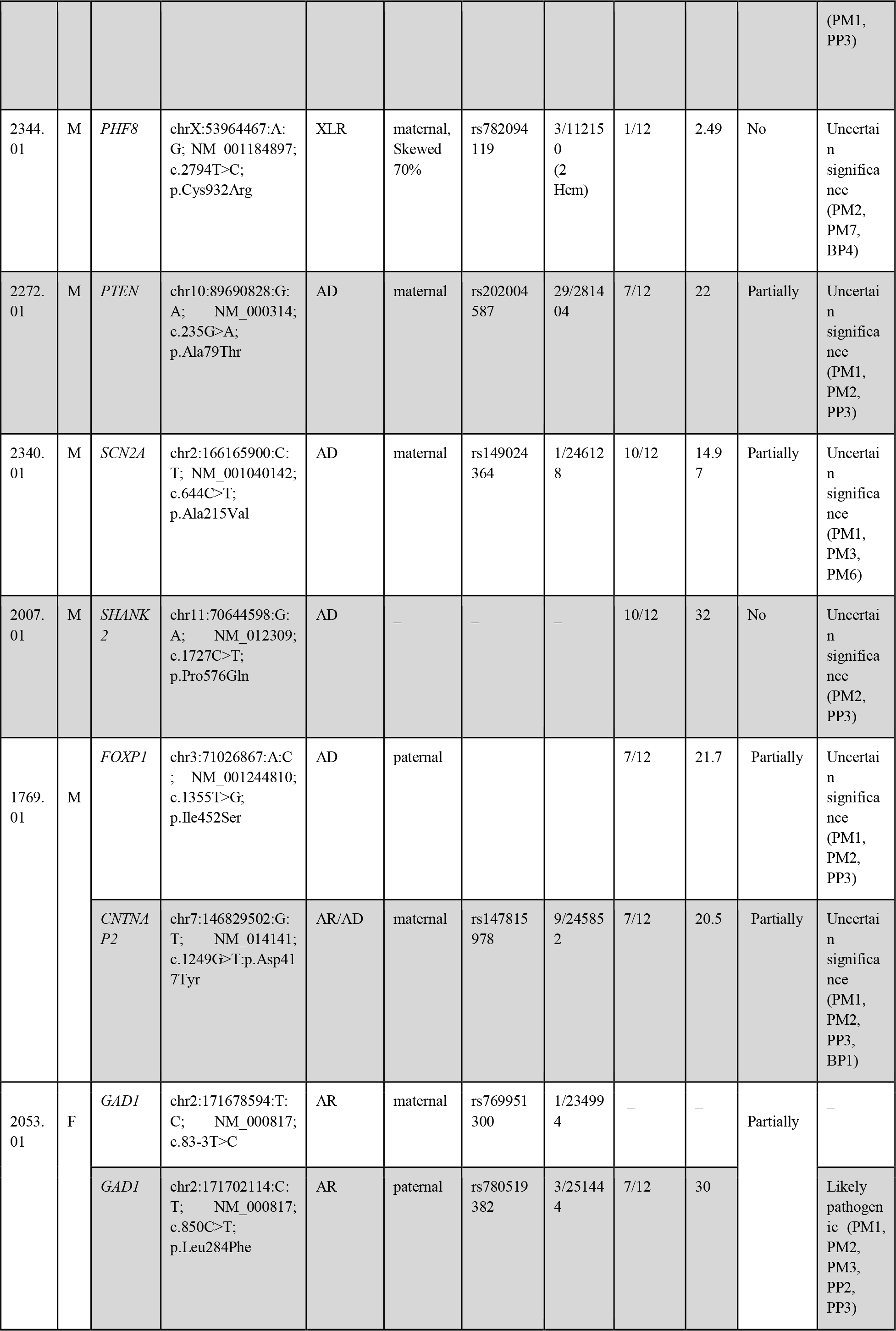
Putative pathogenic variants found in the 150 patients cohort. For maternally inherited variants, the quote of expressed mutated allele in the mother, based on X-inactivation analysis, was reported. Prediction consensus is calculated among the twelve methods provided by ANNOVAR (see Methods section). Variant interpretation based on InterVar: criteria manually adjusted based on our findings (Richards et al., 2015). M, male; F, female; Genomic position on human GRCh37; AD: autosomal dominant; AR: autosomal recessive; XLD: X-linked dominant; XLR: X-linked recessive; Hem: hemizygote.

To support the disease relevance of each selected variant, multiple lines of evidence were required: conservation, allele frequency in population databases, computational inferences of variant effect, mode of inheritance, X-inactivation pattern, and disease segregation.

### *In silico* analysis of candidate variants

Canonical protein sequences were retrieved from UniProt (Apweiler et al., 2004) and protein domains predicted by InterPro (Mitchell et al., 2018). To evaluate conservation, orthologous sequences were downloaded from OMA Browser (Schneider et al., 2007) and aligned with MAFFT (Katoh and Standley, 2013). When available, crystal structures were retrieved from PDB (Rose et al., 2017). Structures of protein domains were modelled with MODELLER (Alva et al., 2016) (automatic best template selection), using templates predicted by HHpred (Alva et al., 2016). Structure of the CASK L27 domain and its complex with SAP97 have been analyzed using Mistic2 (Colell et al., 2018) to evaluate covariation between residues. Structures were manually explored with Pymol (Janson et al., 2017) or UCSF Chimera (Pettersen et al., 2004). Disorder content and the presence of short linear motifs for protein interactions were assessed combining MobiDB (Potenza et al., 2015) and ELM (Gibson et al., 2015), using the interactive exploration tool ProViz (Jehl et al., 2016).

## RESULTS

### Gene panel description

The computational approach adopted to select panel genes includes genes recurrently mutated in ID or ASD conditions, genes shared among ID and ASD disease networks, and genes directly connected to ID/ASD. Of the 74 selected genes, 42 are FMRP targets, 21 postsynaptic proteins, and 16 chromatin modifiers. The majority of the selected genes are associated with autosomal dominand diseases; of these *DEAF1*, *PTEN* and *RELN* are also associated with a recessive disorder. Moreover, the panel includes 8 genes associated with autosomal recessive disorders and 19 genes associated with X-linked diseases. Twenty genes are associated with non-specific ID, while 40 genes are responsible for defined genetic syndromes. Seven genes have been found to confer autism susceptibility (*SHANK2, CNTNAP2, RELN, CHD8, NLGN3, NLGN4X, and PTCHD1*). At the time of panel design, for 11 of the selected genes there was only scant evidence in the literature about their association with ID and/or ASD (*ASH1L, NTNG1, KATNAL2, MIB1, MTF1, MYH10, PTPN4, TANC2, TBR1, TRIO*, and *WAC*). Two more genes associated with an OMIM ID have been meanwhile reclassified and the disease association confirmation is pending (*HDAC4* and *KIRREL3*) (Supplementary Table S2).

### Data Output and Quality

Our strategy allowed generating 263 reads per amplicon with 97% of them on target. The mean target region coverage for all patients ranged between 67 and 612. About 94% of target regions were covered at least 20x and 93% of amplicons had no strand bias. The uniformity of amplicon coverage resulted being 89%. A small proportion of targeted regions was weakly covered (<20x) throughout all patients. These are mainly first exons or GC-rich regions representing a well-known burden in an amplicon based strategy. Specifically, exon 4 of *MECP2*, exon 5 of *ARX*, exon 1 of *FMR1*, part of exon 21 of *SHANK3* have a read depth <10x. Depending on phenotype manifestations, otherwise these regions have been covered by Sanger sequencing, as they could not be analyzed reliably.

### Cohort description and diagnostic yield

Our cohort of 150 individuals is enriched in males (61%). 81.3% were sporadic cases and the remaining had a family history of neurodevelopmental disorders with siblings affected in 3.3% of cases. At the time of molecular testing, the age of the patients was ranging between 2 and 42, with a median of 11 years. Although the vast majority of patients have ID, for four individuals clinicians did not report information about the presence of an intellectual defect. In 38 cases, the level of cognitive impairment has not been evaluated. Among the patients with a cognitive impairment evaluation, a slightly higher proportion have a moderate form (25.3%) than severe (22%) and mild ones (24.6%). 93 patients (62%) had both ID and ASD. 53 (35.3%) had only ID, while 4 (2.6%) had ASD and no information was provided about the presence of ID. Epilepsy was reported for 55 (36.6%) individuals, of which 6 had early onset epilepsy (<24 months). 39 patients with ID and ASD present also epilepsy (26%) (Table 1). MRI and EEG abnormalities were reported, respectively, for 37.3% and 24.6% of the sequenced individuals (Table 1). In most of the patients a structural pathogenic alteration of chromosomes was excluded by FISH, karyotype, or aCGH; however, in 17 of these, aCGH analysis revealed a copy number variations (CNVs) inherited from unaffected parents or involving gene-poor regions (Supplementary Table S3). 86 patients had a negative Fragile-X test, and 57 resulted negative to other single gene tests, such as *MECP2*, *CDKL5*, *UBE3A*, and/or to the Chr15q methylation test.

### Variant detection and prioritization

In coding exons or exon-intron boundaries regions of the 74 genes, we detected on average about 74 single nucleotide variants (SNVs), with a range of 60 to 80 SNVs per patient. Overall, 202 coding or splicing SNVs passing the quality control (GQ>30, DP>20) and frequency filters (MAF<1%), were observed only once in our cohort. Based on the prioritization criteria described in the Methods section, we selected about 170 variants for further analysis, 47 of which were absent from the general population (Supplementary Table S4). We detected certainly causative mutations in 26 patients, leading to an overall diagnostic yield of 17.3% for the entire cohort. These rare or novel variants were predicted pathogenic by several approaches, found to be *de novo* or inherited from affected parents and consistent with the expected disease for the respective gene (Table 2). In other 15 patients, we identified 17 rare variants (missense, synonymous or splicing) with a putative although not established pathogenic role (Table 3). For these variants, either there were no family members available for segregation analysis, or the clinical features did not fit those that would have been expected for the respective gene. With the availability of functional studies or further sequencing data, we expect that a number of these variants will turn out to be benign, but some variants might be proven pathogenic. In particular, five of these putative pathogenic variants meet most of the ACMG criteria (p.Tyr2418Phe in *ATRX*, p.Gly1465Arg in *CREBBP*, c.730+1G>A in *DEAF1*.Val821Leu in *GRIN2B*, p.Pro576Gln in *SHANK2*) and thus with segregation analysis we would be able to assign their causative role on the proband’s phenotype. For two other cases, MR1769.01 and MR2053.01, either the digenic or autosomal recessive transmission is possible, only with functional assays supporting the pathogenic effect of the identified variants. In addition to a novel missense mutation in *FOXP1* gene, MR1769.01 carries a rare variant in *CNTNAP2*, which is transcriptionally regulated by *FOXP1.* As previously proposed by O’Roak and collaborators, we hypothesize a two-hit model for the disease risk in this patient, where a mutant *FOXP1* protein leads to an amplification of the deleterious effects of p.Asp417Tyr in *CNTNAP2* (O’Roak et al., 2011). In MR2053.01, we detected two variants in GAD1 gene, which in associated with an autosomal recessive phenotype. The paternally inherited missense p.Leu284Phe maps on the pyridoxal 5'-phosphate (PLP) transferase domain and is predicted as pathogenic by several computational methods, while the maternally inherited c.83-3T>C is predicted to alter splicing mechanisms. However, we were not able to demonstrate its pathogenicity by qualitative analysis of the transcript extracted from the mother.

After segregation analysis, we were able to classify about one hundred of the selected variants as likely benign. The majority of these variants were found in genes associated with highly penetrant disorders inherited from apparently healthy parents (n=85) or found in individuals with a causative mutation in another gene consistent with the proband phenotype (n=7). Five variants in X-linked genes were inherited from a healthy father or the X-inactivation pattern did not supported their pathogenic role (Supplementary Table S4).

Finally, some rare or novel inherited variants with strong pathogenicity predictions were classified as possible contributing factors. These variants occur on genes known to confer autism susceptibility (e.g. *SHANK2*, *RELN*, *CNTNAP2*) or that have been reported in the literature in individuals with very mild phenotype (e.g. *TRIO*, and *SLC6A1*) (Supplementary Table S4).

### Disease causing variants

We identified likely causative variants in 26 patients (17%) and 18 different genes (Table 2). More than one mutation was found in seven genes (*ANKRD11, CASK, EHMT1, GRIN2B, MECP2, SHANK3*, and *TRIO*); *SHANK3* resulted to be the most mutated gene. Among the identified variants, six were in X-linked genes and 12 in genes associated with autosomal dominant conditions. Twenty variants were absent from the gnomAD database, while five were known pathogenic mutations of the *MECP2, SATB2, SHANK3*, and *GRIN2B* genes.

Most of the mutations were *de novo*; one patient carried two of them. In 20 cases, paternity and maternity were established, while for two others, the parent DNA samples were not available. We can assume that the two truncating variants in *ANKRD11* and *MECP2* should be *de novo*, since they involve genes that are associated with highly penetrant disorders. Furthermore, the p.Arg168X variant in the *MECP2* gene is a recurrent pathogenic variant associated with Rett syndrome. We also identified four inherited causative variants, one maternally inherited variant in the TRIO gene, which is associated with an autosomal dominant disorder, and three maternally inherited variants in X-linked genes (*ATRX, GRIA3*, and *RAB39B*). X-inactivation analysis was consistent with the phenotype expression. The *RAB39B* variant is thought to be also responsible for the mild phenotype reported in the mother, clinically re-evaluated after the molecular finding.

### *In silico* structural analysis may help predict mutation effects

15 causative variants (1 splicing, 6 frameshift, and 8 stop codon variants) are predicted to result in truncated proteins, if escaping the nonsense mediated mRNA decay. The *SHANK3* splicing variant has been shown to functionally impair mRNA splicing, producing an aberrant transcript containing an additional 77 bp intronic sequence (Li et al., 2018). Nine out of the 12 missense mutations were predicted pathogenic by the majority of computational methods, while pathogenicity predictions were discordant for three variants (Table 2). Among the nine variants with strong prediction of pathogenicity, p.Arg696His in *GRIN2B* has previously reported as pathogenic and has been shown to alter the Agonist Binding Domain (ABD) reducing channel activity (Swanger et al., 2016). I*n silico* analysis of two other missense mutations, *DYRK1A* p.Lys174Asn and *EHMT1* p.Gly1193Arg allowed us to classify them as loss of function mutations. Lysine 174 maps to the catalytic pocket of the *DYRK1A* kinase domain and alters the electrostatic surface of the domain, which is important for nucleotide binding (Figure 1). Glycine 1193 maps on the *EHMT1* SET domain, which is necessary for methylation of lysine-9 in the histone H3 N-terminus, and is buried in the rigid structure of the domain. A substitution of Glycine 1193 with an arginine residue should result in unfolding of the domain core (Figure 2).

**Figure 1.**
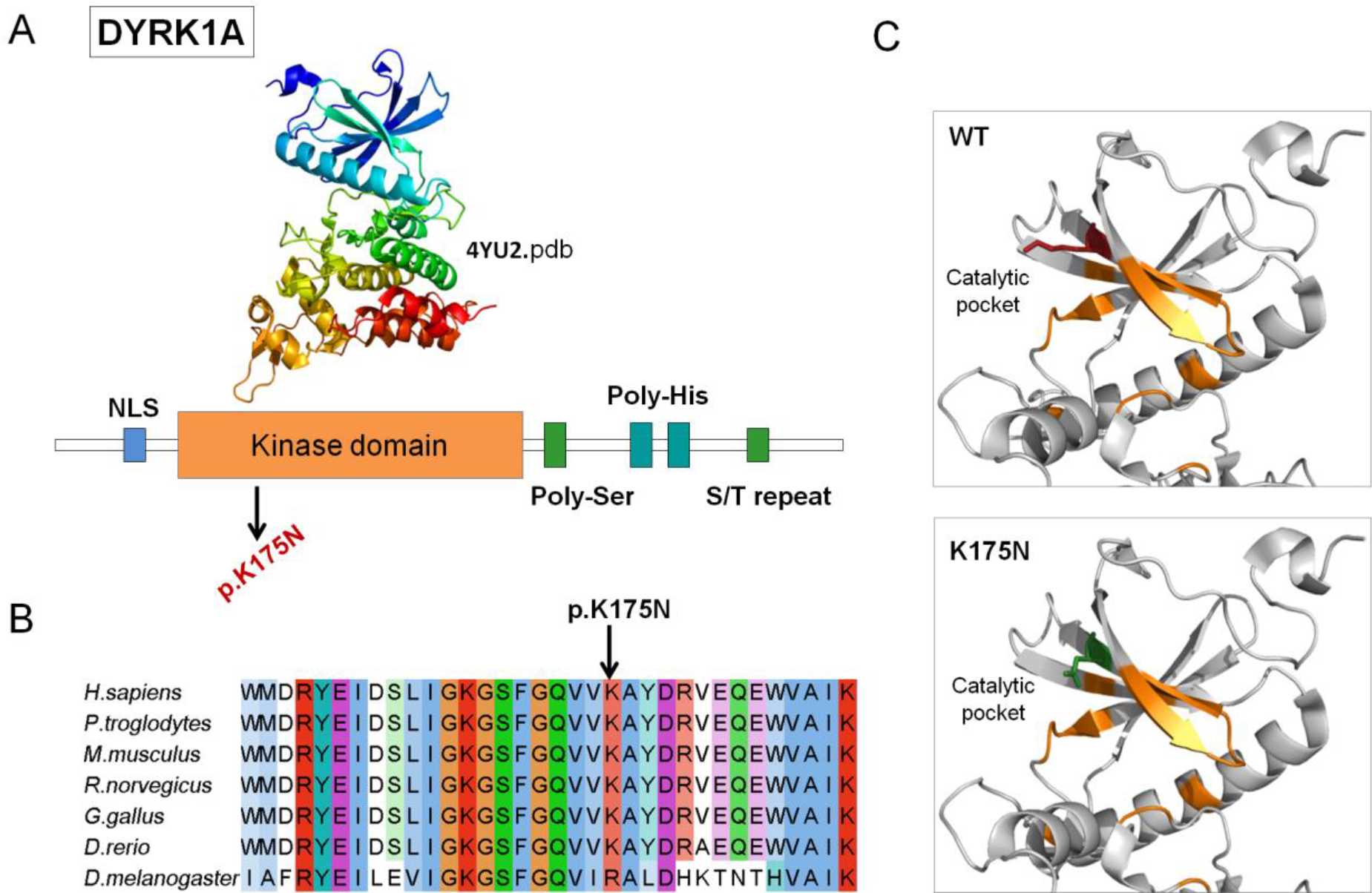
DYRK1A missense mutation affect catalytic pocket of kinase domain. **A)**Domain architecture of DYRK1A. Protein sequence presents regions biased toward polar (serine and threonine) and aromatic (histidine residues). NLS = nuclear localization signal. **B)**DYRK1A p.Lys175 and neighboring residues are conserved among orthologous sequences. Amino acids are colored by conservation, according to ClustalX color code. **C)**DYRK1A p. Lys175Asn variant and wild type residues are mapped to kinase domain structure (4yu2.pdb, chain A). Residues involved in nucleotide binding are represented in orange sticks, wild-type lysine (K) in red and asparagine (N) in green.

**Figure 2.**
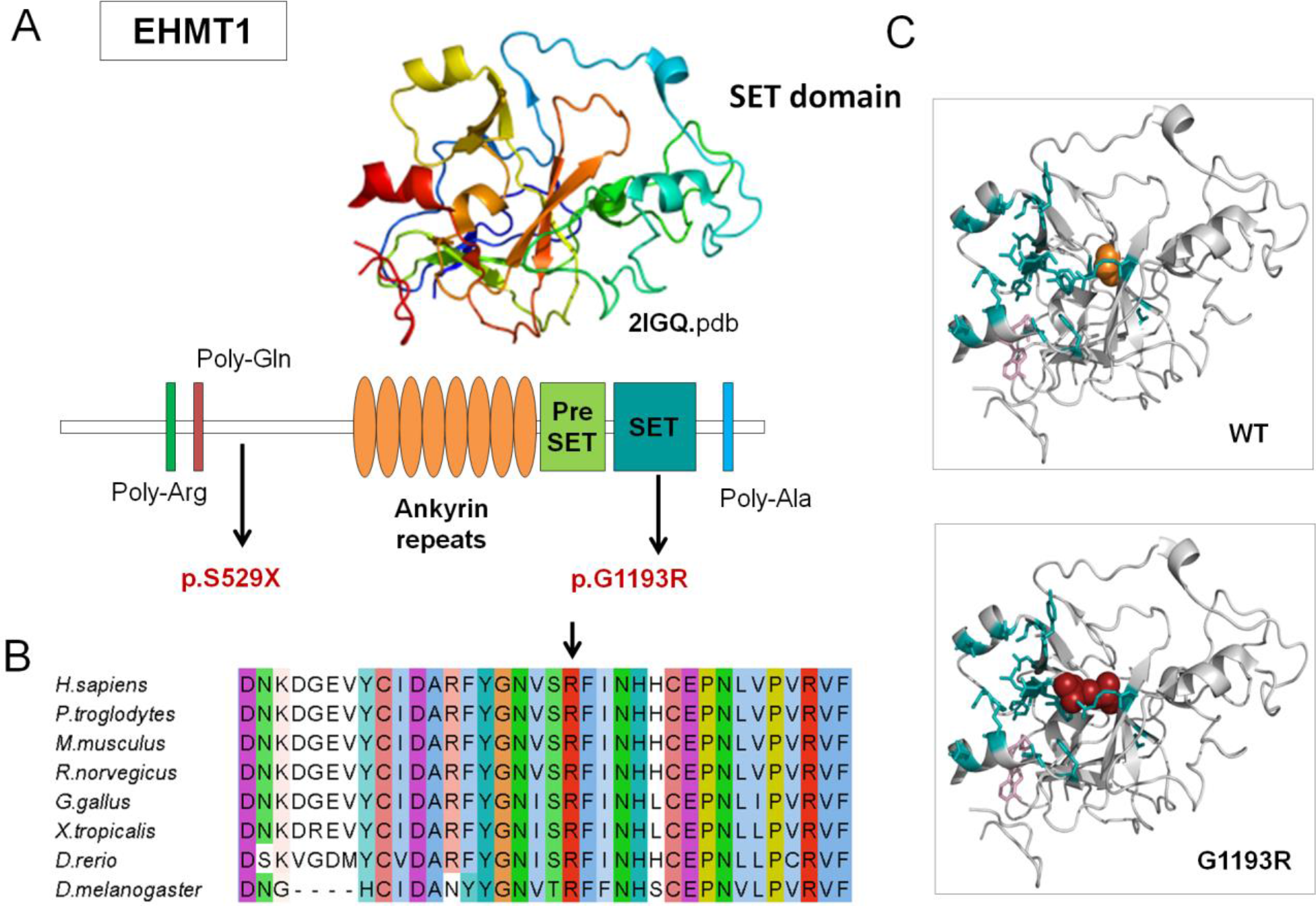
EHMT1 missense mutation affects SET domain. Causative variants identified in our patient cohort are mapped to the EHMT1. Protein sequence presents regions biased toward polar (glutamine and arginine) and a poly-alanine motif. The ankyrin domain (orange) is involved with the histone H3K9me binding. The Pre-SET domain (green) contributes to SET domain stabilization **B)**EHMT1 p.Gly1193 and neighboring residues are conserved among orthologous sequences. Amino acids are colored by conservation, according ClustalX color code. **C)**EHMT1 p.Gly1193Arg variant (red) and wild type glycine (orange) are mapped to SET domain structure (2igq.pdb, chain A) Residues involved in H3K9 binding and the S-adenosyl-L-methionine molecule are represented in blue sticks.

We also hypothesized a predictive impact on the protein function of two variants with discordant pathogenicity: p.Phe193Leu in *RAB39B* (Supplementary Figure S2) and p.Tyr387His in *CASK* (Figure 3). Phenylalanine 193 maps in the RAB39B hypervariable C-terminal tail, which mediates interactions with effector proteins for proper intracellular targeting (Chavrier et al., 1991). This non-conservative substitution may disrupt a functional motif involved in protein interaction and cause a mislocalization of RAB39B, as shown for the protein mutated at position p.Gly192Arg closeby (Mata et al., 2015).

**Figure 3.**
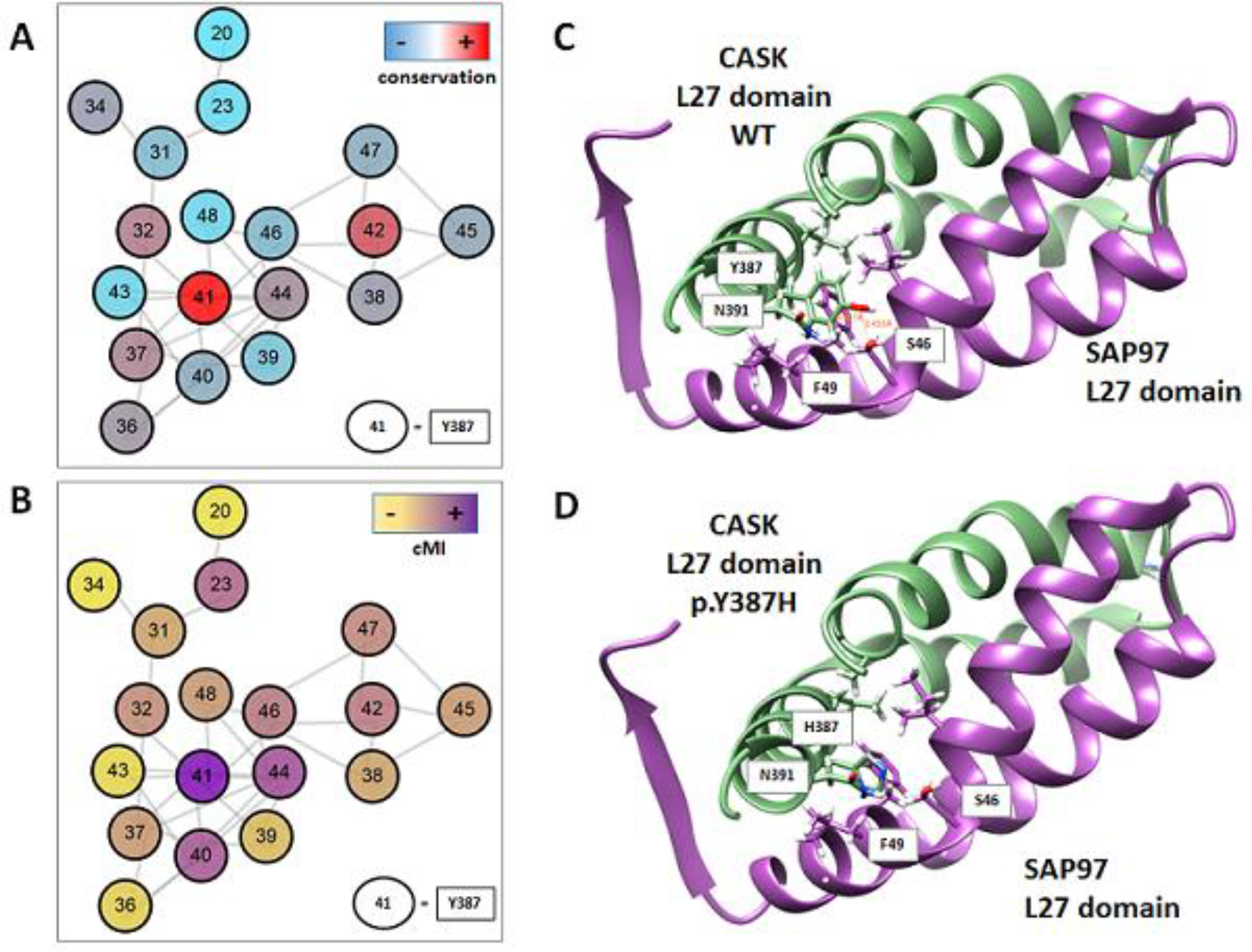
The novel likely hypomorphic mutation of CASK may alter L27 domain dimerization. A) Covariation network of L27 domain (PF02828). Nodes are the residues (L27 domain numbering) coloured by conservation from red to light blue (highest to lowest respectively). B) Nodes are coloured by cumulative Mutual information (cMI) from violet to yellow (highest to lowest respectively). Edges are the top 0.1% covariation scores (Mutual Information) calculated with Mistic2 (Colell et al., 2018). C) Ribbon representation of the L27-SAP97 complex. CASK domain L27 is coloured green (chain B 1RSO PDB) and SAP97 violet (chain A 1RSO PDB). In sticks are shown the residues that interact (their R) with Y387 in the complex. D) same colouring schema. Y387 was “*in silico*” mutated to H387. Mutation and figures were generated by UCSF Chimera (Pettersen et al., 2004).

The Tyr387 residue maps on the N-terminal L27 domain, which mediates hetero- and homo-dimerization of the CASK protein. Comparing L27 domain sequence from different proteins, the 387 position is one of the most conserved and connected in the structure, denoting its important structural role (Figure 3A and 3B). However, at that position histidine is more frequent than tyrosine, thus the p.Tyr387His should not have dramatically effect on the L27 domain structure (Supplementary Figure S3). However, when considering more related homologs of human CASK, the tyrosine 387 is highly conserved, presenting mostly tyrosine or, in few cases, phenylalanine compared to other L27 domains; this might indicate that tyrosine is important for the protein function and that a change to histidine might be deleterious, particularly in human and nearest species. We hypothesize that this substitution may result in a hypomorphic mutant protein with a reduced ability to form dimers (Figure 3), which is consistent with the phenotype of the boy, hemizygous for the *CASK* mutation. Indeed, individuals carrying hypomorphic mutations have been reported with an X-linked intellectual disability with or without nystagmus and additional clinical features (Hackett et al., 2010).

### Genotype – phenotype correlation

One of the major criteria to assign the pathogenicity of the variants is the patient phenotype correlation with the classical syndrome associated to the corresponding gene. Most of the identified causative mutations (n=22) correlate with the previously reported phenotype of the corresponding gene (Table 2). For instance, the two known *MECP2* mutations were found in two girls with a suspected Rett syndrome. The p.Arg168* variant found in MR414.01 patient was missed in previous single gene testing since it was in a mosaic state. The p.Arg294* variant found in MR2145.01 patient was identified by Sanger sequencing of the *MECP2* region not covered by the 74 gene panel. A previous panel for Rett-Angelman spectrum disorders also missed the variant. In addition, for the two cases carrying a *de novo* mutation in *EHMT1* gene, the patient phenotypes were consistent with a Kleefstra Syndrome (KS). Both patients presented with core symptoms of the disease, including a moderate to severe intellectual disability with absent speech, and hypotonia. However, a Kleefstra syndrome was suspected for the characteristic carp-shaped mouth, only in MR2243.01 carrying the frameshift mutation, while MR2166.01 was originally suspected to have Smith Magenis Syndrome (SMS) probably due to brachycephaly, which is also a common characteristic of KS.

Nonetheless, in other cases the probands lacked some peculiar clinical features of the associated syndromes that prevented the geneticists to formulate hypothesis about a suspected syndrome. As an example, the two patients carrying a truncating mutation in the *ANKRD11* gene, both lacked, macrodontia of the upper central incisors, craniofacial features, and skeletal anomalies typical of the KBG syndrome, but both have mild ID, behavioral issues, and hearing loss (Sirmaci et al., 2011).

On the other hand, in some cases the detected mutations were found in genes never associated with particular phenotypic traits. In particular, we observed macrocephaly in three individuals carrying pathogenic mutations in three different genes (*ATRX*, *GRIN2B*, *and TRIO*) previously associated to microcephaly (Table 4). Interestingly, the *de novo* p.Arg696His in *GRIN2B* had been previously reported in a girl with a significant phenotypic overlap with our patient (MR 2019.01), including developmental delay, poor speech, intellectual disability, ASD with stereotypic behavior (Swanger et al., 2016). However, our patient present macrosomia and marked macrocephaly. There is only one report in literature associating *GRIN2B* with macrocephaly, in a case with an ~2 Mb interstitial deletion in 12p13 involving the entire *GRIN2B* gene, in addition to other genes (Morisada et al., 2016). Furthemore, we observed that the other two cases presenting with macrocephaly (MR 2276.01 and MR 984.01), carrying the maternally inherited *ATRX* and the *de novo TRIO* mutations, also carried other rare inherited CNVs or sequence variants (Supplementary Tables S3 and S4). Since inherited from healthy parents these alterations were classified as benign or of uncertain significance.

**Table 4:** Mutated genes in the different phenotypic manifestations (ASD, epilepsy, Microcephaly, Macrocephaly, Hypotonia, and Ataxia) Some genes have been found mutated in individuals presenting phenotypic traits that have not been previously associated to these genes (highlighted in red, bold).

| Clinical features | N° individuals affected (Tot=150) | Genes carrying causative variants | Genes carrying likely pathogenic variants |
| --- | --- | --- | --- |
| Intellectual disability |  |  |  |
| Autistic traits | 73 | ANKRD11, ARID1B, CASK, EHMT1, GRIA3, GRIN2B, MECP2, MED13L (ASH1L), SHANK3, SYNGAP1 | ANKRD11, CHD8, FOXP1 (CNTNAP2), DEAF1, GRIN2B, KATNAL2, PHF8, PTEN, SCN2A |
| Epilepsy | 40 | ANKRD11, EHMT1, GRIA3, MECP2, OPHN1, RAB39B, SETBP1, SYNGAP1, TRIO | ANKRD11, CASK, CHD8, CREBBP, GAD1, SCN2A, SHANK2 |
| Microcephaly | 19 | ARID1B, CASK, DYRK1A, MECP2, TRIO | **CHD8**, CREBBP, **SHANK2** |
| Macrocephaly | 12 | **ATRX**, GRIA3, **GRIN2B**, **TRIO** | - |
| Hypotonia | 28 | CASK, DYRK1A, EHMT1, MECP2, **TRIO** | - |
| Ataxia | 11 | MECP2 | GAD1 |

## Discussion

An accurate clinical and molecular diagnosis can greatly improve the treatment of individuals with neurodevelopmental disorders. However, differentiating between pathophysiological conditions that are clinically and genetically heterogeneous is a big challenge particularly when they arise with common comorbid disorders. The recent advent of Next Generation Sequencing (NGS) technologies allowed discovering many genes involved in these conditions. The study of undiagnosed cases with similar clinical manifestations through whole exome or genome sequencing highlighted the wide spectrum of phenotypic expression of some well-known genetic conditions, such as Rett syndrome, that can be caused by different genes involved in common biological pathways (Ehrhart et al., 2018). We are moving from a phenotypic-based to a gene-centered view of the diseases and we are incorporating the concept of disease network in the classification of the genetic conditions (Barabási et al., 2011). Due to the cost, ethical problems, and storage resources limitations, the use of genome and exome sequencing in the clinical practice is still a challenge. However, the availability of bench top systems, such as Ion Torrent platform, allowed spreading the use of gene panel sequencing in diagnostic laboratory for testing different genes involved in heterogeneous conditions, such as neurodevelopmental disorders.

In this study, we show the application of targeted sequencing of 74 genes in a cohort of 150 individuals with ID and/or ASD designed for diagnostic purpose. A certain causative variants have been found in 17.3% of the tested individuals, the diagnostic yield is the same in the 93 individuals presenting ID and ASD comorbidity. In contrast to other studies, the majority of the diagnosed cases have a more severe ID (Redin et al., 2014) (Table 1). Considering patients presenting other comorbidities, such as epilepsy, microcephaly or macrocephaly, the diagnostic yield further increases to 20%, 31.6%, and 36.4%, respectively. Thus, the ability to find a molecular cause with this selection of genes is more effective in severe cases than those with a mild phenotype are. The use of network parameters to filter panel genes allowed selection of core genes, which are often hub genes implicated in multiple biological processes. Thus, an alteration of these genes may have consequences in diverse neurological systems leading to a more severe phenotype.

Interestingly, in three patients presenting macrocephaly as one of the clinical manifestations, we found a causative variant in three different genes (*ATRX*, *GRIN2B*, and *TRIO*) that have not been associated before to this feature. We decided to classify these variants as potentially causative even if the proband’s phenotypes were only partially consistent with the phenotype reported for each gene. In these cases, to support the pathogenic role of the variants we considered the complex genetic architecture underlying the pathogenic mechanisms involved in NDDs (Woodbury-Smith and Scherer, 2018). The p.Arg696His in *GRIN2B* gene has been previously reported in an individual with ID but no macrocephaly, and experimentally tested for its ability to impair the protein function (Swanger et al., 2016). The other two cases carrying a *de novo TRIO* and a maternally inherited *ATRX* mutation were found to carry other rare inherited SNVs or CNVs. We can speculate that these rare inherited alterations may contribute to the disease as modifier variants interacting with causative ones to determine a specific phenotype. Diverse rare CNV, such as 15q11.2 and 16p12.1 deletions, have been implicated in multiple neurodevelopmental disorders, and a multi-hit model have been suggested (Girirajan et al., 2010; Abdelmoity et al., 2012). Furthermore, a multifactorial model has been proposed to explain the heritability of ASD, where rare *de novo* and inherited variations act within the context of a common-variant genetic load (Chaste et al., 2017; Guo et al., 2018)). This might also explain the variable clinical outcomes associated with the same causal variant, such as in the case of p.Arg696His in *GRIN2B*.

Another consideration to take into account is that for some NDD candidate genes a clear description of the related disorder will become possible only with the accumulation of reported cases. The *TRIO* gene has been recently associated with mild to borderline intellectual disability, delay in acquisition of motor and language skills, and neurobehavioral problems. Other findings can include microcephaly, variable digital and dental abnormalities, and suggestive facial features (Ba et al., 2016; Pengelly et al., 2016). Only few individuals carrying pathogenic mutations in this gene have been reported to date, thus the clinical manifestations of the *TRIO*-related disorder is still evolving. Here, we report two novel individuals carrying pathogenic *TRIO* mutations, expanding the phenotypic spectrum associated with this novel NDDs gene. Furthermore, different mutations in the same gene can have different effects on the gene product, and therefore different pathological consequences (Barabási et al., 2011). These variants may perturbe specific subset of links in the interactome. For instance, different mutations in *ARX* gene, a paradigm of a pleiotropic gene, have been associated with diverse defects involving GABAergic neurons and associated with a wide spectrum of disorders (Friocourt and Parnavelas, 2010). This may also be the case for the p.His1371Tyr mutation in the GEF1 domain of *TRIO*. Recently, it has been show that variants affecting different protein interaction interfaces of this domain can produce bidirectional alterations of glutamatergic synapse function. The p.His1371Tyr maps far away from the Rac1 binding interface, near another variant, the p.Asp1368Val, that has been previously reported in a boy with severe ID (de Ligt et al., 2012). In contrast to other GEF1 domain variants involved in ASD, the p.Asp1368Val has been demonstrated to results in TRIO hyperfunction (Sadybekov et al., 2017). This finding may explain the more severe phenotype associated to the p.Asp1368Val, and those of the proband we report, carrying the p.His1371Tyr, presenting with severe ID, epilepsy, absent speech, and macrocephaly.

With the targeted approach, a proportion of patients remains without molecular diagnosis due to the limited number of sequenced genes. However, the higher coverage obtained by the gene panel sequencing, compared to a whole genome or exome sequencing approach, allows detecting a high number of variants in the target regions. Furthermore, due to the relatively small number of variants to be further investigate, this approach allows focusing on rare variants that would be filtered out for discordant pathogenicity predictions, as the novel hypomorphic *de novo CASK* mutation found in hemizygous state in a male with intellectual disability. For these variants, an in depth *in silico* analysis of the protein structure and function allowed to pinpoint possible mutation effects to support their pathogenic role. Finally, with this approach we provided a set of genotype-phenotype associations for a core set of genes involved in NDDs, which can be used to train or test computational methods for prioritization of potential disease-causing variants.

### CAGI-competition

At the completion of our study and before submitting for publication, genetic data and corresponding phenotypes of the tested individuals have been provided for a challenge at the Critical Assessment of Genome Interpretation (CAGI). CAGI is a worldwide blind test to assess computational methods for predicting phenotypic impacts of genomic variations. The outcome of the predictive competition using the ID-ASD gene panel data set are presented elsewhere. However, here we provide the original data with few updates that can be used to train and/or to test their own computational tools/approach aiming to predict comorbid phenotypes from genetic variants in a subset of NDDs genes.

## Conclusions

The heterogeneity of NDDs reflects perturbations of the complex intra- and intercellular networks. The emerging tools of genomic medicine allows to hold the promise of disentangling the complex genetic architecture of these particular disorders, leading to the identification of different etiologies in similar phenotypes as well as common pathways underlying apparently distinct conditions. Based on the finding that diseases that share genes or involve proteins interacting with each other show elevated comorbidity, we designed a 74-gene panel to perform sequencing of individuals with ID and ASD comorbidities. With this approach, we identified disease causing variants or putative pathogenic variants in 27% of the tested individuals. This work demonstrates that knowledge about shared genes and common pathways can be used to develop innovative diagnostic tools needed to discriminate among overlapping phenotypes with high risk of developing comorbid features.

## Acknowledgments

The authors are grateful to all the probands families and clinical Institutions that referred the patients to the Laboratory of Molecular Genetics of NDDs, as well as to members of the BioComputingUP group and Molecular Genetics of NDDs, for insightful discussions. This work was supported by Italian Ministry of Health Young Investigator Grant GR-2011-02347754 to E.L and S.C.E.T.; Fondazione Istituto di Ricerca Pediatrica - Città della Speranza, Grant 18-04 to E.L. Data reported in this work were used as challenge at the CAGI-5 organized by S. E. Brenner, J. Moult, and Gaia Andreoletti, and titled “Predict patients’ clinical descriptions and pathogenic variants from gene panel sequences”. “The CAGI experiment coordination is supported by NIH U41 HG007446 and the CAGI conference by NIH R13 HG006650.

## Disclosure statement

The authors declare no conflict of interest.

