## Supplementary figures and images for "Characterization of Intellectual disability and Autism comorbidity through gene panel sequencing"

### Supplementary Figure S1

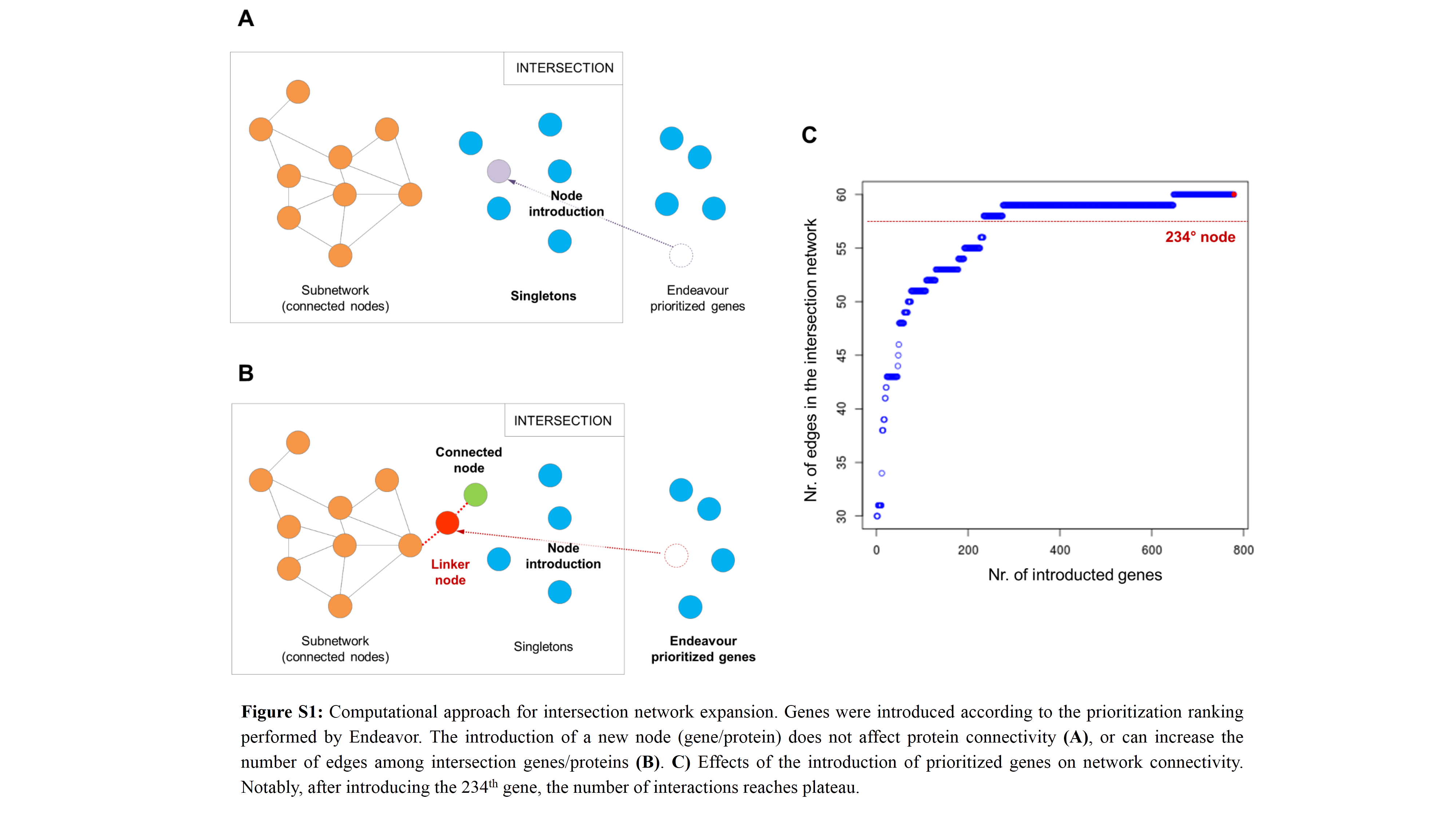

### Supplementary Figure S2

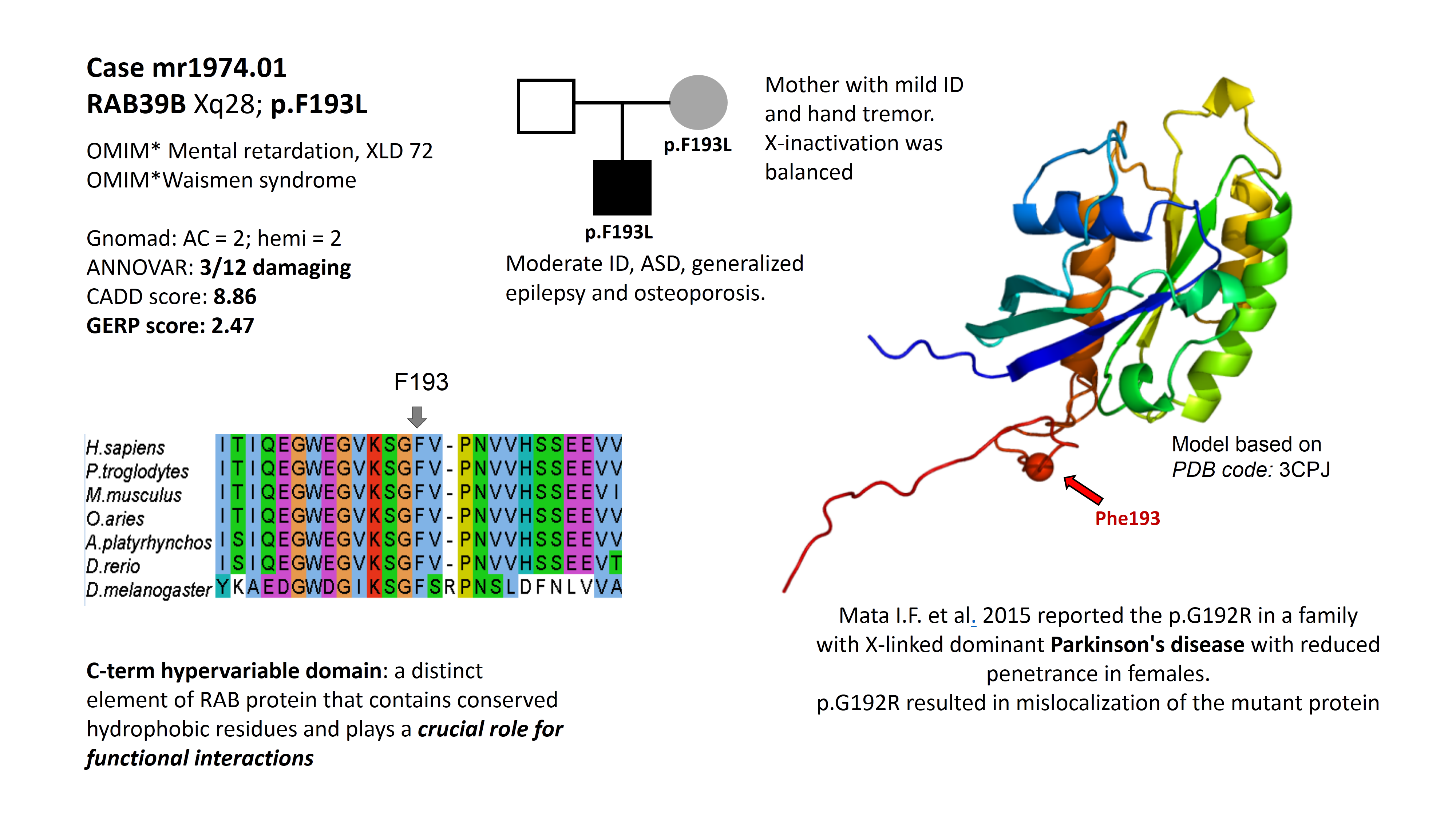

### Supplementary Figure S3

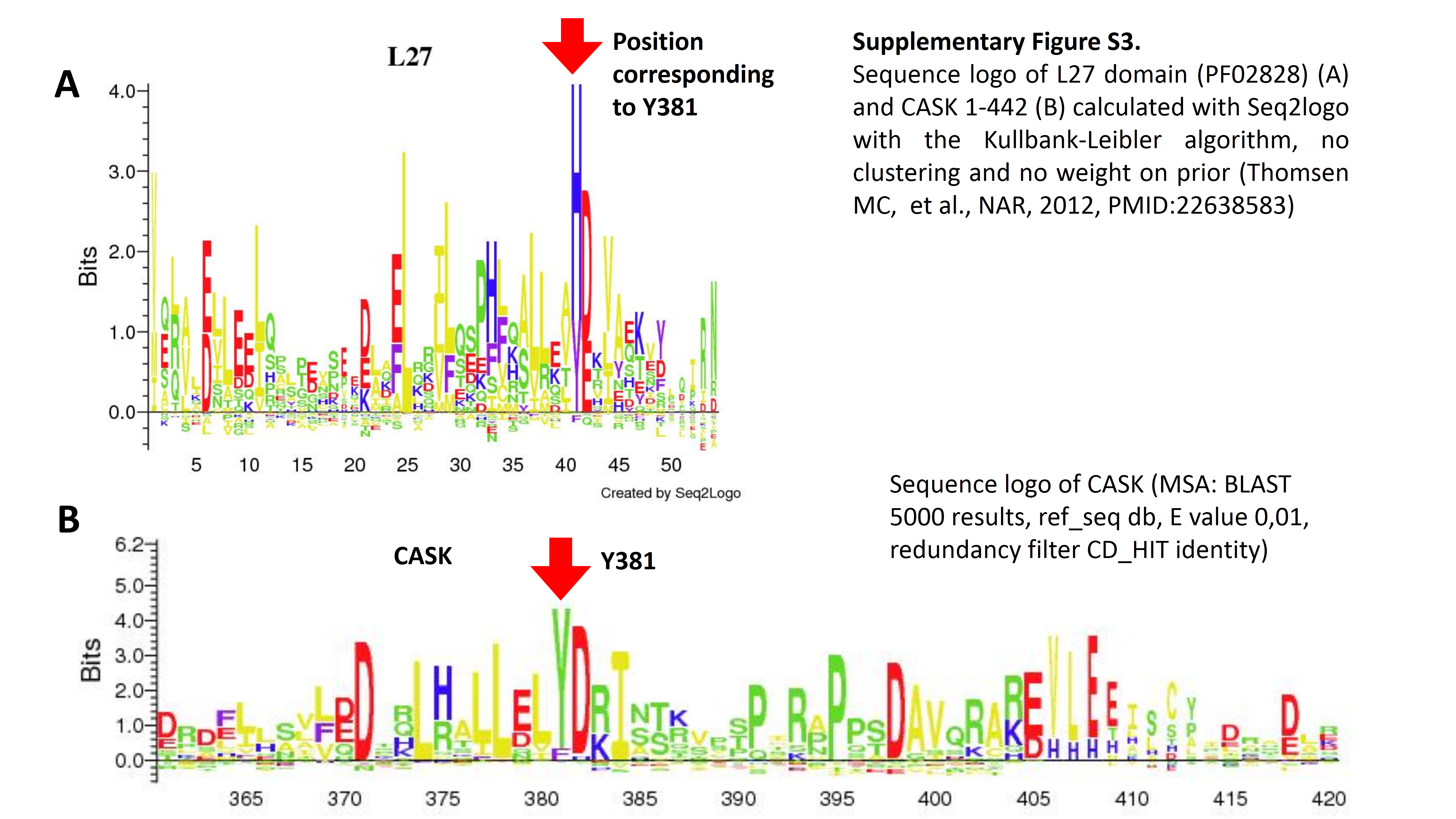
