## Supplementary Table S1 for "Characterization of Intellectual disability and Autism comorbidity through gene panel sequencing"

| **Reference for known ASD/ID associated genes** | |
| --- | --- |
| **Phenotype** | **Source** |
| ASD | Table S3: ASD candidate genes, only those with at least 2 references and list from (2) (1) |
| ASD | Table S5: ASD genes (3) |
| ASD | Table S6A: ASD genes (4) |
| ID | Table S6: ID genes (3) |
| ID | Table S6C: ID genes (4) |
| **Reference for candidate ASD/ID genes from whole exome/panel sequencing** | |
| **Phenotype** | **Source** |
| ASD | Table 2: Summary of confirmed de novo events (5) |
| ASD | Table 1: Top de novo ASD risk contributing mutations (5) |
| ASD | Table 2: Loss of function in probands (6) |
| ASD | Table 3: Likely disrupting mutations in affected children (7) |
| ASD | Table S1: All validated de novo events and annotations (no synonymous variants) (3) |
| ASD | Table 1: Summary ASD genes found with mutations in this study (8) |
| ASD | Supporting Table 5: Genes with Damaging, Validated Variants in More Than One Family (1) |
| ASD | Supplementary Table 3: The list of de novo variants identified in 48 ASD cases (only frameshift, nonsynonymous, stop gain) (9) |
| ASD | Table 2: Gene disrupting rare variants shared by affected sibs in each multiplex family (10) |
| ID | Table 3: Missense, nonsense, frameshift, and splice site de-novo variants in genes associated with intellectual disability in each patient–parent trio (11) |
| ID | Table 4: Probable disease-causing de-novo variants in each patient–parent trio (11) |
| ID | Table 2: Genes Affected by De Novo Mutations Associated with Intellectual Disability (12) |
| ID | Table 2: Overview of all de novo variants identified by exome sequencing in ten individuals with unexplained mental retardation (13) |
| ID | Table 2: Top risk DNMs identified in this study (14) |
| ID | Table 2: List of all causative/possibly causative mutations identified in our cohort (15) |
| ID | Tables 2-4: **Homozygous and compound heterozygous variants validation using Sanger sequencing and in silico prediction** (16) |

**Supplementary Table S1. Sources used to collect genes associated to ID or ASD.**
