## Supplementary Table S3 for "Characterization of Intellectual disability and Autism comorbidity through gene panel sequencing"

| Patient code | Sex | Alteration | Chr position | Size | Inheritance |
| --- | --- | --- | --- | --- | --- |
| MR984.01 | M | dup | 12q24.32 | 153,5kb (interstitial) | paternal |
| MR1730.01 | F | dup | Xq26.2 | . | maternal |
| MR1769.01 | M | dup | Xq23 | 221kb | maternal |
| MR1861.01 | F | dup | 4q28.2 | 135,3kb | maternal |
| MR1999.01 | F | dup | dupXq21.33 | 379 Kb (region with few genes) | . |
| MR2023.01 | M | dup | 15q11.2 | 1Mb (Between *BP1* e *BP2*: including *TUBGCP5*, *CYFIP1*, *NIPA1*, *NIPA2*) | maternal |
| MR2039.01 | M | dup | 8p22 | . | paternal |
| MR2072.01 | F | dup; dup | 10p11.21; 16p13.2 | 275kb; 487kb | maternal; maternal |
| MR2142.01 | M | del; del | Yq11.22; Yq11.23 | 1,9Mb; 1.4Mb | paternal; paternal |
| MR2201.01 | M | del | 6p25.2 | 105kb (including *SLC22A23*) | . |
| MR2222.01 | M | dup | 2q37.7 | of uncertain significance | . |
| MR2239.01 | F | del | Xq27.2 | . | maternal |
| MR2276.01 | M | del | del1q31.1 | 188Kb | maternal |
| MR2318.01 | F | dup | dup4p35.2 | 778.9Kb | maternal |
| MR2340.01 | M | dup; dup | 11p13; Xq21.2 | . | paternal; maternal |
| MR2347.01 | M | dup | 9q33.3 | . | maternal |
| MR2349.01 | M | dup | 11p14.3 | gene-free region | . |

**Supplementary Table S3.** CNV alterations found in patients of the cohort. Most of the variants have been considered as benign or of uncertain significance.
